# Unusually Broad-spectrum small-molecule sensing using a single protein scaffold

**DOI:** 10.1101/2025.05.15.654352

**Authors:** H Tian, J Beltrán, W George, C Lenert-Mondou, N Seder, ZI Davis, SD Swift, T Girke, TA Whitehead, I Wheeldon, SR Cutler

## Abstract

Small-molecule sensing in plants is dominated by chemical-induced dimerization modules. In the abscisic acid (ABA) system, allosteric receptors recruit phosphatase effectors and achieve nM *in vivo* responses from µM receptor–ligand interactions. This sensitivity amplification could enable ABA receptors to serve as generic scaffolds for designing small-molecule sensors. To test this, we screened collections of mutant ABA-receptors against 2,726 drugs and other ligands and identified 569 sensors for 6.7% of these ligands. The mutational patterns indicate strong selection for ligand-specific binding pockets. We used these data to develop a sensor design pipeline and isolated sensors for multiple plant natural products, 2,4,6-trinitrotoluene (TNT), and “forever” per- and polyfluoroalkyl substances (PFAS). Thus, the ABA sensor system enables design and isolation of small-molecule sensors with broad chemical scope and antibody-like simplicity.

## Main text

The ability to create protein binders through antibodies and designed proteins has transformed biotechnology research by making essentially any protein a targetable entity (*1*). The success of this strategy is partly due to the simplicity of using a single protein domain and engineering pipelines for isolating binders. Developing protein scaffolds that can be programmed to recognize small molecules as easily as antibodies can recognize antigens would empower many biotechnology and synthetic biology applications. In principle, numerous biological parts can be reprogrammed to develop small-molecule sensors, including ligand-induced transcription factors (for example, *lacI*) (*2*), cell surface receptors (for example, GPCRs and DREADDs) (*3*), and chemical-induced dimerization (CID) systems (for example, rapamycin/FKBP/FRB) (*4*). Of these, CID systems are particularly attractive because they enable the facile control of many outputs via induced-proximity mechanisms (*4*–*8*).

Genetic dissection of phytohormone signaling in land plants has uncovered many naturally occurring CID modules that could potentially be harnessed to engineer new sense-response capacities (*9*–*13*). We recently introduced the plant ABA receptor PYR1 (*Pyrabactin resistance 1*) as an engineerable scaffold for sensor design and demonstrated the ability to reprogram binding for several cannabinoid and pesticide ligands (*14*–*17*). Part of the success of the PYR1 receptor as a scaffold is due to the separation of ligand-recognition and binding from its coreceptor, HAB1 (*Homolog of ABA Insensitive 1*). This simplifies engineering efforts by restricting designs to a single molecular recognition surface. PYR1 also benefits from intrinsic sensitivity amplification. The native ligand, ABA, binds to the receptor with micromolar affinity, but the activated receptor conformer binds with nanomolar affinity to HAB1 in an orientation that blocks ligand dissociation, leading to an up to 100-fold increase in affinity to the ligand (*18*). This amplification of sensitivity reduces the challenges of identifying alternative activating ligands that have low affinity (*19*). Moreover, its components are soluble and function in diverse prokaryotic and eukaryotic hosts (*12, 14, 16, 20*). These features coalesce to make the PYR1 CID module an effective sensor scaffold and a candidate for designing small-molecule binders akin to antibodies for protein antigens.

Here, we set out to define the chemical scope of the PYR1 binding pocket and its potential to be used as an antibody-like scaffold for sensing small molecules. To do this, we established a miniaturized sensor-isolation pipeline that capitalizes on ligand-induced growth triggered by the physical interaction between PYR1 and HAB1 using yeast growth-based selections (**Figure 1**). Using this approach in thousands of parallelized assays, we screened receptor variants against a collection of 2726 small molecules. Together, these yielded sensors for 182 molecules, 6.7% of the ligands screened. We show that this large dataset of ligand-receptor interactions can be harnessed to isolate high-affinity sensors in a single step with antibody-like simplicity.

**Figure 1.**
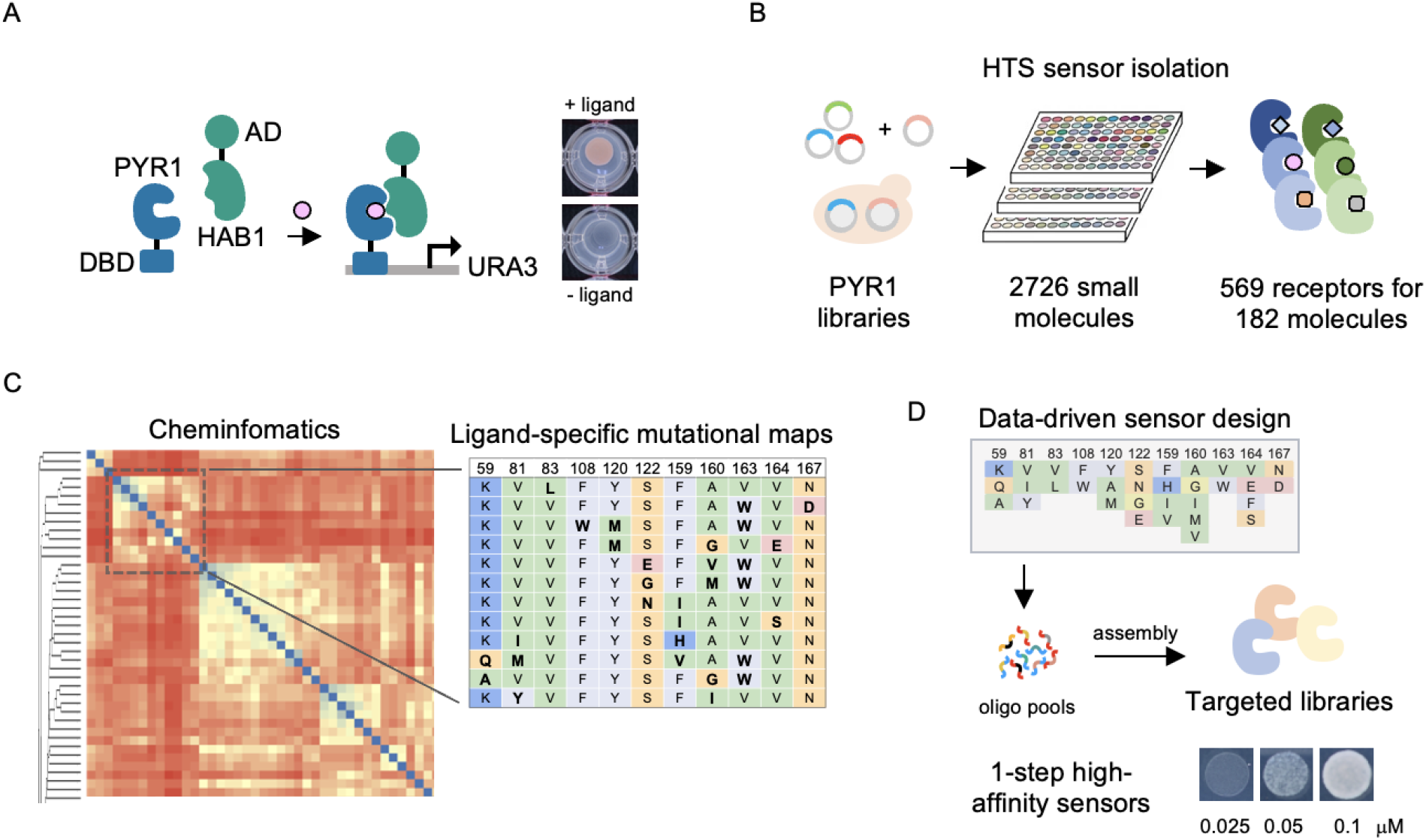
Large-scale sensor isolation to enable data-driven sensor design. (**A**) The PYR1 sensor isolation pipeline. PYR1 binds to its effector protein HAB1 in response to ABA, its native ligand. This interaction can be measured in yeast growth assays by activating a genetic circuit that rescues uracil auxotrophy. Mutant PYR1 variants that recognize new ligands are identified from receptor library pools using selection experiments (responders grow without uracil). (**B**) A high-throughput version of the receptor isolation approach was developed. In this, we plate ~150,000 cells onto media containing a test chemical and select colonies from the wells for retesting and subsequent sequencing. These efforts identified 569 receptors that recognize 182 unique molecules. (**C**) Chemoinformatic analysis of the data is used to build maps of sequence-ligand interactions. (**D**) The sequence-ligand maps developed can be harnessed for data-driven design of sensors. Libraries of receptors targeted to specific molecule classes are constructed using sequence profiles to design oligonucleotide pools that can be assembled using Golden Gate cloning; subsequent growth-based screens enable one-step isolation of high-affinity sensors.

To profile the binding scope of PYR1, we assembled a collection of diverse small molecules from a commercially available collection of FDA-approved drugs and natural products, and a curated set of agrochemicals and other research ligands (**Table S1)**. Since commercial screening libraries often contain redundancy, for example, the same parent molecule present as different salts, we used ChemmineR (*21, 22*) to define the non-redundant set of molecules screened. We mapped them to the ChEMBL database (*23*) to identify redundancies and extract molecular properties and approval statuses. These analyses showed that the screening collection contains 2726 small molecules. This collection was screened in duplicate against previously and newly designed PYR1 variant libraries harboring double and triple mutations localized to 19 ligand-contacting residues (see **Methods** for details) (*14*). Candidate sensor strains for a given molecule were screened in duplicate at 100 µM and retested at multiple concentrations of the molecule to eliminate false positives and profile the sensitivity of the strain. Together, these screens yielded 569 distinct sensor sequences and strains that respond to 182 small molecules (**Table S2**), including 80 FDA-approved ligands, 31 plant natural products, 17 agrochemicals, and 54 research molecules, representing an overall success rate of 6.7% of the ligands screened (**Figure 2A**).

**Figure 2.**
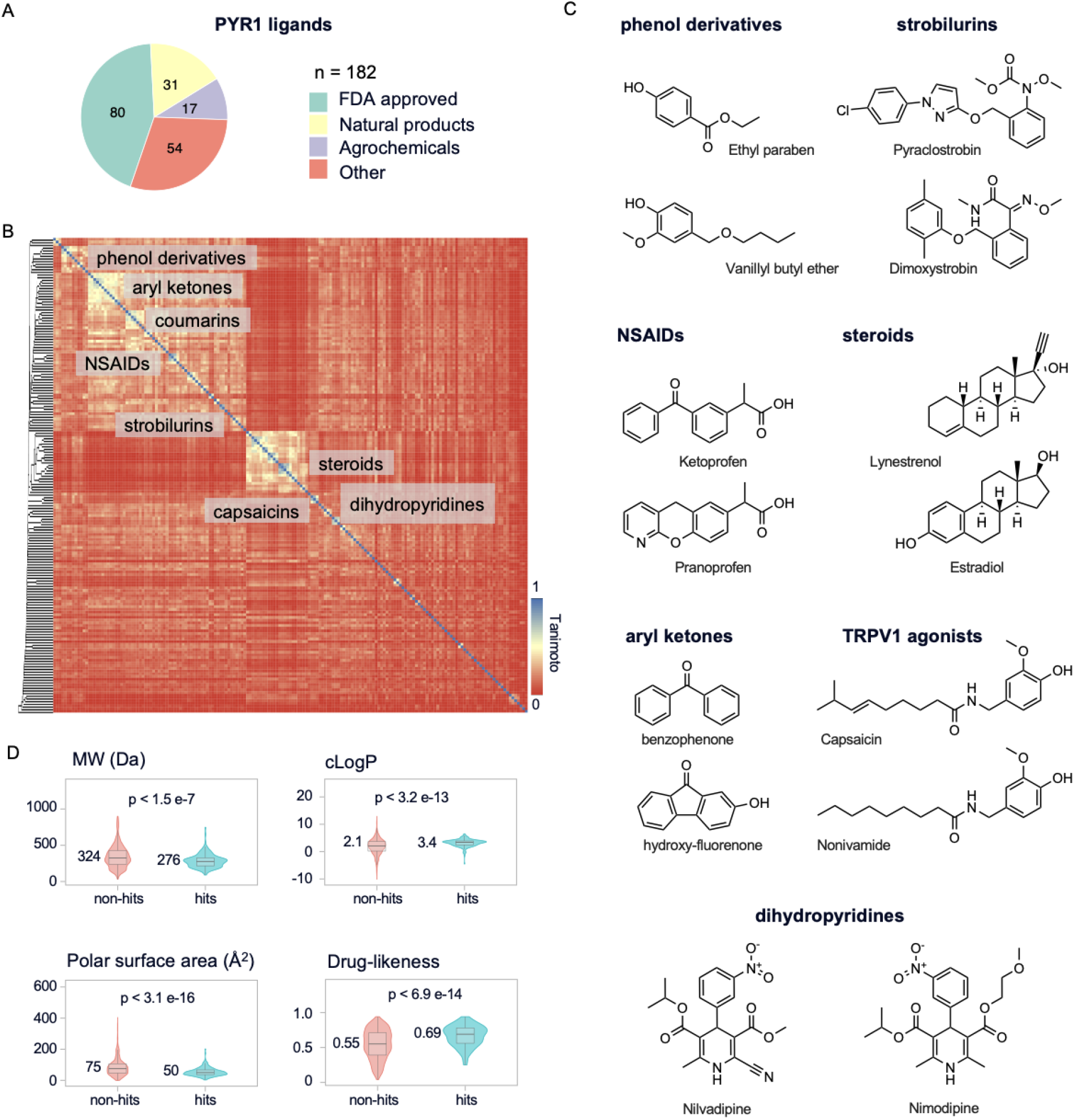
The PYR1 binding pocket can be mutated to sense small molecules of broad chemical scope. (**A**) Summary of ligands sensed by receptors isolated from a 2726-member small molecule library screen. The hits obtained include FDA-approved molecules, natural products, agrochemicals, and a collection of research ligands (see **Table S1** for the ligands screened). (**B**) The molecules sensed by PYR1-derived sensors are structurally diverse. The heatmap shows hierarchical clustering based on the pairwise chemical similarities of the 182 hit molecules. Pairwise ligand-similarity,, measured using Tanimoto scores calculated from atom-pair descriptors in ChemmineR (*21*), is shown by the heatmap (blue = identical). Selected clusters of similar ligands are highlighted. (**C**). Selected chemical ligands from each cluster identified in part B. (**D**) PYR1-derived receptors sense drug-like molecules. Comparisons of the physicochemical properties of ligand hits (n=182) and the non-hits (n=2544). Molecular weight (MW), computationally calculated octanol-water partition coefficient (cLogP), topological polar surface area, and a quantitative estimate of drug-likeness (*24*) values were extracted from ChEMBL (*23*). The median of each population is shown, along with the p-value from a two-sided Wilcoxon rank sum test.

Chemical clustering of the hit ligands based on their pairwise Tanimoto distances, calculated using atom-pair fingerprints, reveals a broad distribution of structurally dissimilar hits, with several small clusters of similar molecules (**Figure 2B, C**). Thus, there is broad structural diversity amongst the collection of sensed ligands. Several important pharmacological agents and agrochemicals are sensed by the collection, including steroids, non-steroidal anti-inflammatory drugs (NSAIDs), dihydropyridine calcium channel antagonists, Transient Receptor Potential Vanilloid 1 (TRPV1) receptor agonists, and antifungal strobilurins (**Figure 2C)**. The physicochemical properties of the 182 ligands sensed are drug-like by multiple measures. 171 (96%) comply with Lipinski’s rule of 5, compared to 77% for the library. Moreover, they score highly using the quantitative estimate of drug-likeness (QED) metric, a unitless measure of drug-likeness (med. QED_hits_ = 0.65, QED_non-hits_ = 0.55) (*24*). Compared to the non-hit members of the collection screened, the ligands sensed are enriched for higher cLogP (computed log_10_octanol/water partition coefficient) and lower tPSA (topological polar surface area; **Figure 2D**). These trends are likely driven partly by the cell-based nature of our sensor isolation method, which enriches for molecules with good membrane permeability and bioavailability. The size of ligands sensed is relatively smaller than those of the non-hit collection (median MW_hits_= 274 Da; MW_non-hits_= 324 Da), but includes molecules as small as 90 Da and as large as 748 Da (**Figure 2D**). Collectively, these data demonstrate that the chemical space accessible with PYR1-derived sensors is structurally diverse and drug-like, which defines PYR1 as an unusually versatile and malleable scaffold for developing sensors for drugs and other bioactive ligands.

Analyses of the PYR1-ligand binding pocket dataset show that only 5% of the receptors recognized three or more ligands, suggesting that relatively few low-selectivity receptors were obtained in the set (**Figure 3A**). Consistent with this observation, the mutations obtained are distributed broadly across the ligand-binding pocket (**Figure 3B**). However, F159, which resides at the top of the pocket and contacts both the ligand and HAB1, was preferentially mutated and may be important for stabilizing PYR1-HAB1 interactions. The mutational patterns obtained also suggest strong selection for specific ligand-receptor interactions (**Figure 3C** and **Table S2**). For example, six sensors for the anticonvulsant valproic acid were isolated, and all harbor the mutation I110R, suggesting our screens selected for a stabilizing salt bridge between valproic acid’s carboxylate and the mutant receptors’ basic arginine side chains. Similarly, 11 receptors were identified for the amine-containing ligand cinacalcet, used to treat hyperthyroidism, and 10 of these harbored K59D/E, suggesting selection for a stabilizing salt bridge. These PYR1 sensors suggest a strong selection bias toward distinct binding pocket mutations (**Figure 3B**).

**Figure 3.**
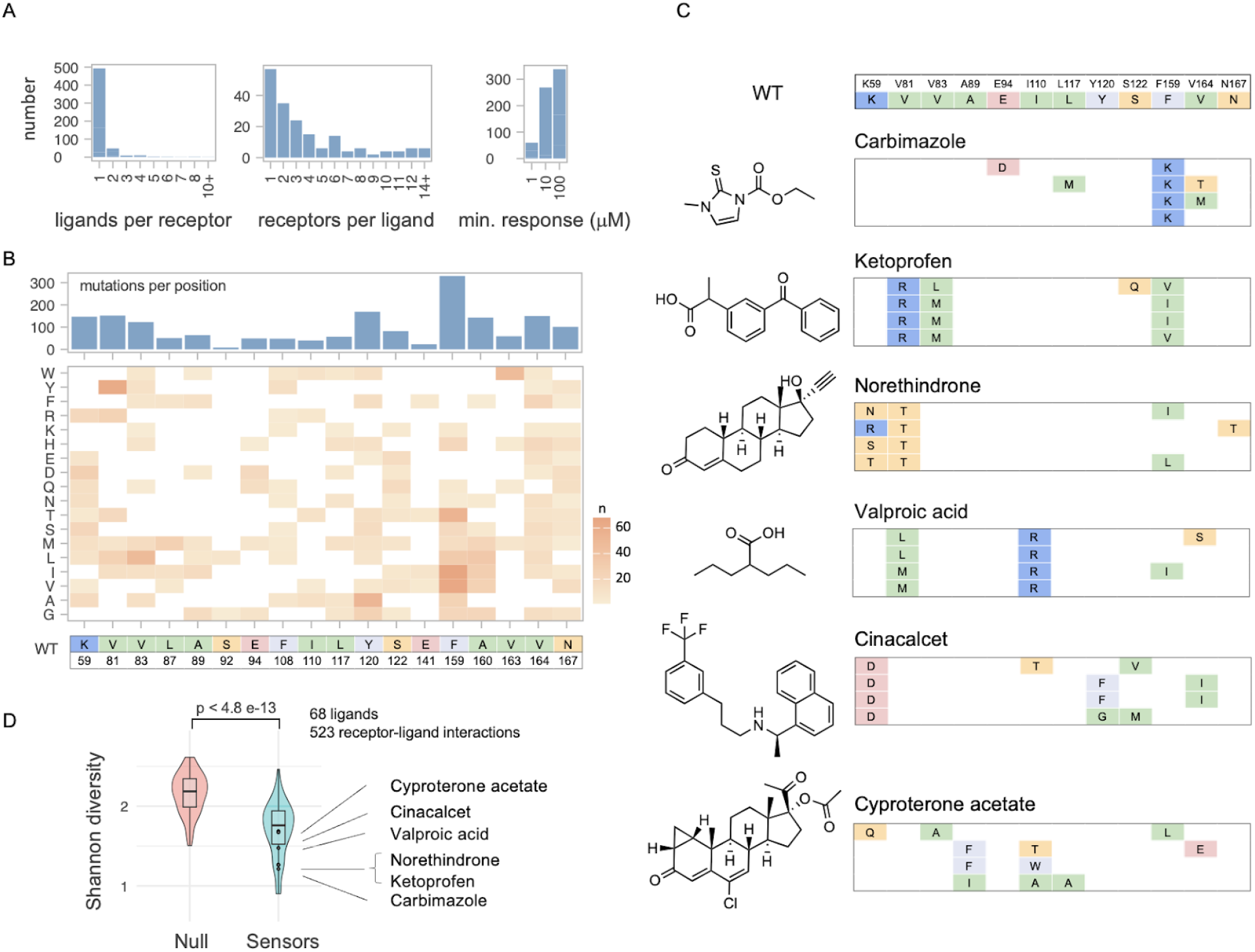
Mutant PYR1 receptor-ligand interactions can be mapped using large-scale sensor screens. (**A**) Screening outcomes: ligand hits per receptor sequence, number of receptors identified per hit ligand, and the minimum dose response of receptor hits during validation (each sensor strain was tested for ligand-dependent growth on 1, 10, 100 µM ligand). (**B**) Sensor mutations are broadly distributed throughout the PYR1 binding pocket; the top panel shows mutations per binding pocket residue, and the bottom panel maps the frequency of specific mutations per residue for the 569 receptors isolated. (**C**) Selected binding-pocket sequences for ligands that yielded ≥ 4 receptor hits. WT residues are shown at the top; colored boxes denote basic (blue), hydrophobic (green), polar (orange), and aromatic (grey) substitutions. (**D**) Sensor binding pocket mutations are tailored to their target ligands. Selection for mutational bias in each sensor-ligand set was quantified by calculating the Shannon diversity of mutation counts per position across receptors, with lower values indicating more positionally clustered mutational patterns (i.e., sensors for a ligand tend to have mutations in the same binding pocket locations). The analysis was performed for the 68 ligands with four or more sensors isolated (n=523 receptor-ligand interactions). The null model shows Shannon diversity values calculated from randomly permuted receptor–ligand assignments (p < 2.5 × 10^−13^; two-sided Wilcoxon rank-sum test).

To quantify these patterns, we employed Shannon diversity to measure the distribution of mutations across binding pocket positions using a shuffled dataset as a null comparator (**Figure 3D**). The lower Shannon diversity values observed in the sensor hits indicate significantly clustered, positionally biased mutational patterns for specific ligands (p < 2.5 × 10^−13^). Lastly, yeast surface display assays demonstrated nM to µM limits of detection for a subset of receptors that functioned well in the YSD assay (**Figure S1**). These experiments demonstrate direct physical interactions without a cell-based growth assay. Collectively, these analyses show strong global selection for ligand-specific mutational patterns, with mutations broadly distributed across the binding pocket.

Our large dataset of ligand–receptor interactions provides maps of binding-pocket residues that, in principle, could be harnessed for receptor improvement. We took two complementary approaches to test this. First, we mined our data for the coumarin scaffold—the largest cluster of plant-derived natural products in our screening library (selected examples shown in **Figure 4A**). Of the 14 coumarins present in the library, we isolated sensors for eight, which harbored mutations at 11 of the 19 binding pocket sites mutagenized. We derived a coumarin-binder sequence profile from these data and encoded combinations of up to nine of the most frequent coumarin-associated mutations (**Figure 4B, Table S3**). The library contained 77,327 members and was constructed by the combinatorial assembly of 218 oligonucleotides (**Table S4**). It was then transformed into *S. cerevisiae* and screened against 12 coumarins (**Table S5**). This yielded sensors for three ligands not identified in the original screens (**5, 6**, and 5,7-dihydroxy-4-methylcoumarin), and between 10-to 400-fold sensitivity gains for the eight positive coumarin hits, including (**1**), (**2**), and (**3**) (**Figure 4C, Figure S2, Table S6**). As a second approach, we asked whether profiles derived from receptors that recognize single-ring phenyl derivatives (**6**) **-** (**9**) could guide the discovery of sensors for the chemically similar molecules, TNT (**11**) and three of its degradation products (**12, 13**, and 4-amino-2,6-dinitrotoluene) (**Figure 4D, S3**); neither of these were present in the screening collection and DSM/TSM library screens did not yield hits for TNT. A 506,229-member receptor library built using a sequence profile derived from receptors activated by (**6**)-(**9**) yielded initial sensors for the four targets (**Table S3**). Sequence profiles derived from the first-round sensors were used to design a second-round library, which ultimately yielded sensors for TNT and three degradation products, DNT and 2ADNT, and 4ADNT with low µM EC_50_ ligand responsiveness, as measured using reporter and growth assays (**Figure 4F, S3; Table S7**). Together, these data demonstrate that our large dataset of ligand-PYR1 interactions can be harnessed to yield receptors with improved affinity and target recognition.

**Figure 4.**
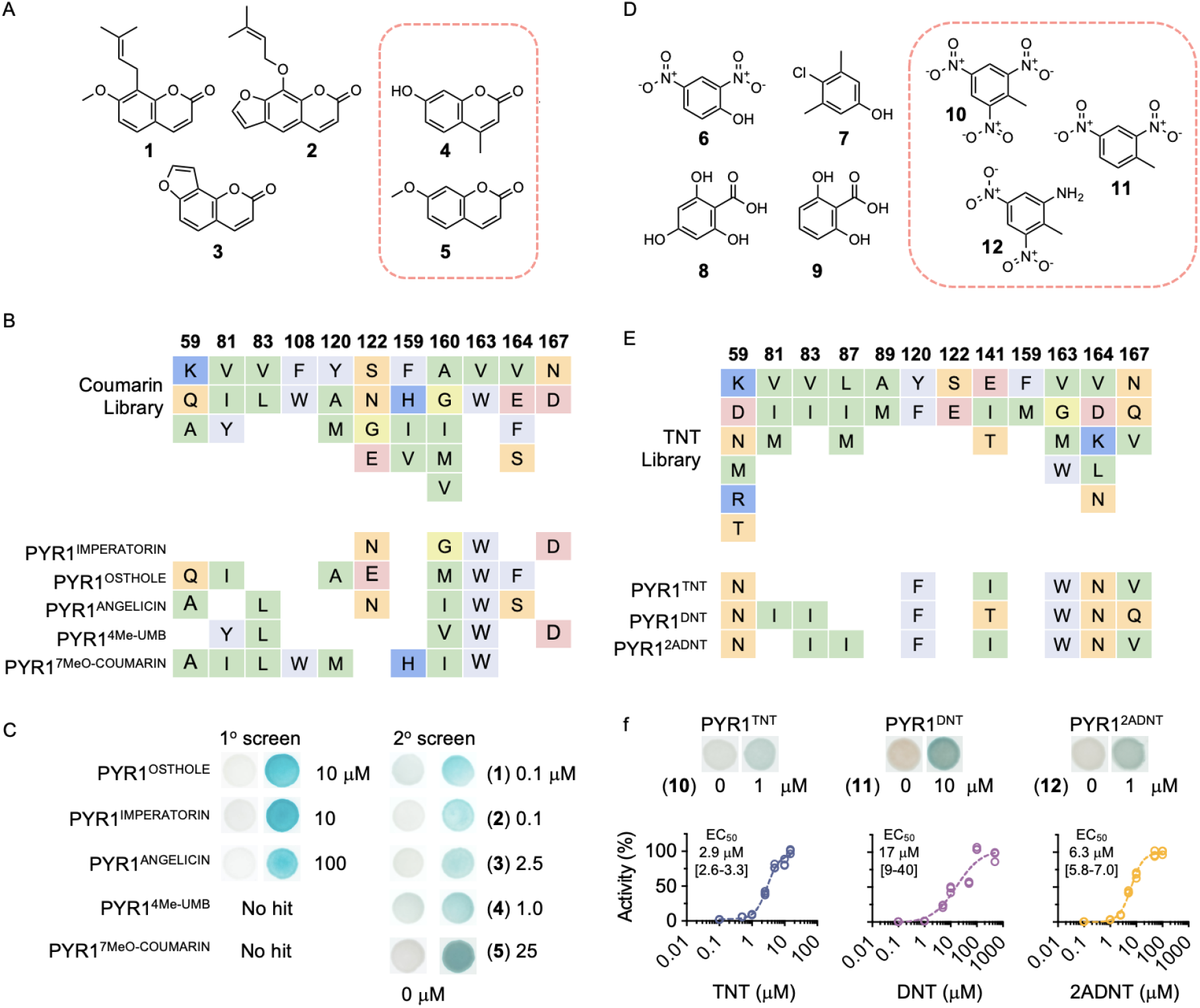
Ligand-receptor interaction maps and sequence profile-guided design enable the isolation of high-affinity sensors. (**A**) Selected chemical structures of coumarin hits used to generate the sequence profile of the coumarin-target library: imperatorin (**1**), osthole (**2**), and angelicin (**3**). Receptors for 4-methylumbelliferone (**4**) and 7-methoxycoumarin (**5**) were not identified in the initial screen; sensors for these compounds were isolated from the coumarin-target library. (**B**) Sequence profile of the coumarin targeted-library and binding pocket mutations of PYR1 receptors activated by compounds (**1**-**5)**. (**C**) Yeast-hybrid β-galactosidase colony assays comparing PYR1 sensor response of the initial sensor and high-affinity sensors isolated from the coumarin-targeted library. The targeted library yielded PYR1 receptors for (**4**) and (**5**), which were not identified in the initial data generation screen. The minimum dose responses are shown in comparison to the mock condition. Figure S2 shows additional growth and reporter dose response assays for the coumarin sensors. (**D**) Chemical structures of selected ligands used to design the sequence profile of a TNT target library: 2,4-dinitrophenol (**6**), chloroxylenol (**7**), 2,4,6-trihydroxybenzoic acid (**8**), 2,6-dihydroxybenzoic acid (**9**). Hits for trinitrotoluene (TNT; **10**), dinitrotoluene (DNT; **11**), and 2-amino-dinitrotoluene (2ADNT; **12**) were not identified using the screening libraries. Sensors for these were isolated from the TNT target library. (**E**) Sequence profile of the TNT target library (TNTv2) and binding pocket mutations for TNT, DNT, and 2ADNT receptors isolated from the library. The sequence profile used for library construction additionally includes 117L/W and 160A/V (**Table S3**). (**F**) Yeast-hybrid β-galactosidase colony and cell-based titration assays for PYR1^TNT^, PYR1^DNT^, and PYR1^2ADNT^ for their on-target ligands. Triplicate data points are shown. The inset data indicates the EC_50_ and range.

Our data establish a simple approach for creating new sensors for small molecules: 1) screen receptor libraries with relatively few binding pocket mutations to identify initial sensors, 2) use the data generated to derive sequence profiles, and then 3) design and screen affinity maturation libraries to yield high-affinity sensors.

To test this pipeline on a class of molecules dissimilar from members of our initial screening library, we focused on ‘forever’ PFAS molecules, which are critical targets for biosensing due to their persistence, bioaccumulation, and associated health and environmental risks (*25*). To do this, we constructed and screened an improved double-site mutant library designed to enhance coverage in HTS assays by reducing auto-activating and wild-type receptors from the original DSM library (DSM-Hao; see **Methods**). We screened this against a panel of 103 different PFAS (**Table S5**), including six of major concern that have been targeted for regulation in drinking water by the EPA (*26*), perfluorononanoic acid (PFNA; **13**), perfluorooctanesulfonic acid (PFOS; **14**), perfluorooctanoic acid (PFOA; **15**), perfluorohexanesulfonic acid (PFHxS; **16**), hexafluoropropylene oxide dimer acid (HFPO-DA), and perfluorobutanesulfonic acid (PFBS) (**Figure 5A**). The initial library screens yielded 86 sensors for 18 targets, providing a dataset to create a PFAS-targeted library for isolating high-affinity sensors (**Figure 5B; Table S8**). Screens using this targeted library expanded the scope of PFAS sensing, increasing the total number of PFAS molecules sensed to 25, and yielded sensors with improved on-target sensitivity. PYR1^PFAS^, isolated in a screen against PFOA (**13**), shows a strong response to low µM concentrations of four of the six molecules targeted for regulation (**Figure 5D, S4**) and provides a new sensor for the most problematic environmental “forever” molecules. These experiments further validate the data-driven sensor approach and its power for developing new sensors.

**Figure 5.**
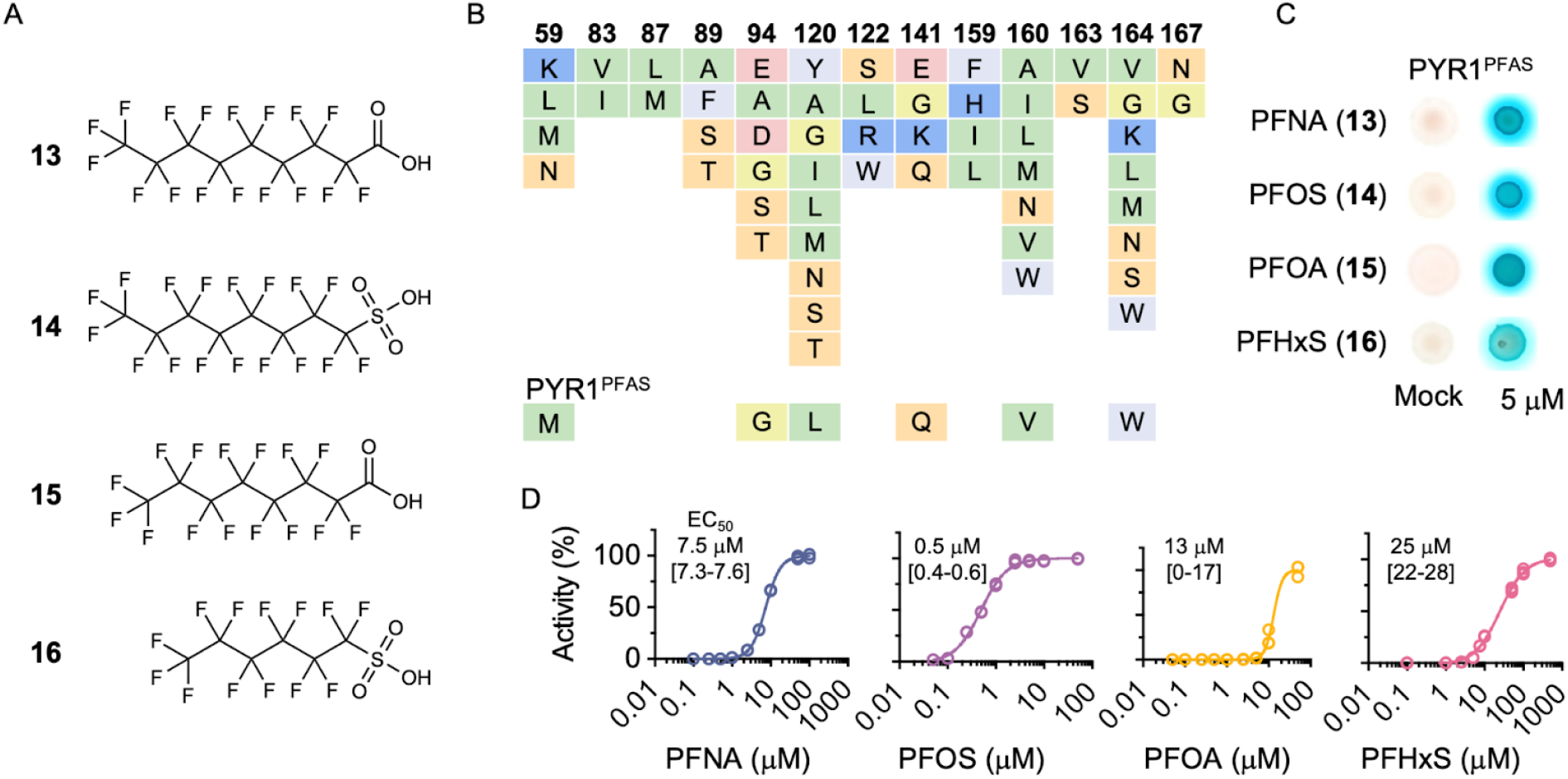
Data-driven engineering of PYR1 receptors for PFAS detection. (**A**) Chemical structures of selected PFAS molecules targeted for sensor design. PFNA (**13**), PFOS (**14**), PFOA (**15**), and PFHxS (**16**) are four of six PFAS molecules regulated by the U.S. Environmental Protection Agency in drinking water. (**B**) PFAS-library sequence profile and binding pocket mutations in PYR1^PFAS^, a pan-PFAS receptor strongly activated by any of (**13** - **16**). The library was screened against 103 unique PFAS molecules (see **Table S5**). (**C**) Yeast-hybrid β-galactosidase colony assays show that PFNA, PFOS, PFOA, and PFHxS strongly activate PYR1^PFAS^. (**D**) Dose-dependent response of PYR1^PFAS^ to four on-target PFAS molecules in cell-based β-galactosidase assays. Triplicate data points are shown. EC_50_ values and range are shown as insets.

## Discussion

The ligand-binding pocket of PYR1 is unusually malleable; it can be reshaped to recognize a surprising number of structurally unrelated ligands. We isolated sensors for 6.7% of the nearly 3000 molecules screened and isolated over a dozen more using data-driven targeted libraries. It has been estimated that drug-like space encompasses approximately 10^33^ molecules (*27*). Even without improvements in hit rates, which should be possible, our system provides access to a large chemical space that can be harnessed to build new biochemical, cellular, and whole-organism control systems. Our sensors recognize drugs and drug-like molecules, opening many possibilities. For example, NSAID-controlled switches, of which we isolated several, could be valuable in clinical applications. Our sensors can also be orthogonalized for deployment in plants using the PYR1*-HAB1* module, which enables two-channel control systems and control of plant phenotypes without perturbing native ABA signaling (*16*). Thus, PYR1’s extreme malleability, ease of sensor design, and induced dimerization mechanism converge to make PYR1 a privileged scaffold for sensor design across biological kingdoms.

Why is PYR1 imbued with such unusual malleability? PYR1 belongs to the ligand-binding SRPBCC superfamily (SRPBCC: START/RHO_alpha_C/PITP/Bet_v1/CoxG/CalC). These are characterised by a helix-grip motif, which consists of a seven-stranded antiparallel β-sheet that wraps around a C-terminal α-helix to create a central ligand-binding pocket. SRPBCC proteins are present in all domains of life and, like our PYR1 variants, bind structurally diverse molecules – from hydrophobic steroids to polar flavonoid-glucosides (*28*–*30*). This domain is frequently involved in transport and catalysis. However, PYR1 and its relatives link ligand-recognition to effector-binding to control signaling. PYR1, therefore, couples an evolutionarily malleable binding pocket with a biochemical mechanism well-suited for engineering signaling systems. Given the success of computational design for CID sensors (*8, 31*) and new methods built for PYR1 (see Leonard *et al*.), we anticipate that PYR1’s properties can be harnessed broadly to design sense-response systems controlled by user-specified molecules with antibody-like simplicity.

Plant hormone signaling employs a suite of sensor modules that, like PYR1, involve hormone-mediated stabilization of protein-protein interactions (*13, 32*–*35*). For example, the gibberellic acid receptor is a member of the large GDSL lipase family and, like the ABA system, its module employs induced dimerization with an effector that greatly stabilizes the activated receptor complex (*36*–*38*). GID1 and other plant hormone sensors should provide a suite of scaffolds for building new sensor systems with the antibody-like simplicity and scope of the PYR1 system.

## Acknowledgments

We thank David Nelson for comments and suggestions on the manuscript.

## Author contributions

Conceptualization: SRC & IW

Methodology: JB, HT, WG, TG

Investigation: JB, HT, WG, CLM, NS, ZD, SS

Supervision: SRC, IW, TAW

Visualization: SRC, IW, TAW, HT, WG, CLM, ZD, SS

Funding acquisition: SRC, IW, TAW

Writing - original draft: SRC, IW

Writing - reviewing and editing: SRC, IW, TAW, JB, HT, WG, CLM, TG, ZD, NS, SS

## Data and materials availability

The main text and supporting methods provide GitHub links to the code used in the manuscript. All chemicals tested, sensor sequences, and oligonucleotide sequences are provided as supplementary tables. Benchling links to the vector sequences used are included in the supplementary materials.

## Materials and Methods

### Chemicals & Libraries

The ligands screened included a commercially available collection of FDA-approved molecules, natural products, and other medicinal compounds (Selleck USA, L1300), which we supplemented with 480 agrochemicals and other ligands purchased from Sigma-Aldrich (USA) and Microsource Discovery Systems (USA). Ligands were purchased as 10 mM stocks in DMSO (**Table S1)**. Screens focused on identifying optimized sensors for specific compound subsets (coumarins, TNT, PFAS) were purchased from various sources as analytic grade standards; **Table S5** lists the chemicals used and their sources.

### Chemical informatics

Compounds were mapped to ChEMBL(*23*) using CAS IDs and compound names, respectively, obtained from the compound library providers to retrieve community identifiers, annotations, and physicochemical and structural descriptors. Due to inconsistent or ambiguous use of CAS IDs and compound names, a multi-step mapping strategy had to be employed. First, Selleck compounds were mapped to DrugBank (*35*) via CAS IDs, followed by ChEBI (*36*) and PubChem. Ambiguous mappings were then resolved using fingerprint/atompair similarities from ChemmineR. CAS IDs obtained from DrugBank, ChEBI, and PubChem were subsequently mapped to ChEMBL. Functional annotations and physicochemical properties were then gathered from various databases, including ChEMBL, ChemmineR (*21*), CDK (*37*), and PubChem. Finally, parent molecule hierarchies (representing the core structure of a compound) from ChEMBL were incorporated into a comprehensive master table. LATCA, an internal UCR collection of curated bioactives from which we selected agrochemicals and other molecules, lacked CAS IDs; these compounds were associated with ChEMBL using their given names and then annotated using a comparable method. The code employed for these annotations can be found in this GitHub repository: https://github.com/tgirke/Map_CMPs_to_ChEMBL.

### Receptor Screening Libraries

Three screens against the collection of compounds were conducted using the same general procedure but with different PYR1 variant libraries. The previously described ~37,797-member library of PYR1 single and double site mutant library (DSM) (*14*) was used in an initial screen and led to the identification of 285 sensors for 113 ligands; a second “shuffle” library containing 1.1 × 10^6^ clones was created using nucleotide excision and exchange technology (NexT)(*38*) to shuffle plasmid DNA for the initial 285 hit sensors; creating a combinatorial library that we screened to yield an additional 23 sensors for 10 ligands. The binding pocket sequences of the hits from the DSM and shuffle sensors were analyzed and used to inform the design of a profile-based library that made all possible ~324,000 triple mutant combinations of the input mutations; **Table S3** shows the sequence profile, and **Table S9** the library coverage statistics (see below for assembly methods). The libraries used were transformed into the *S. cerevisiae* reverse two-hybrid strain MaV99 (*39*) harboring pACT-HAB1, which expresses a HAB1-activation domain fusion that enables ligand-mediated activation of a *UAS*_*GAL*_::*URA3* expression construct and rescue of MaV99’s uracil auxotrophy. Transformations were conducted using the LiAc/SS carrier DNA/PEG method (*40*) to generate ≥11x library coverage. The yeast libraries were then subjected to two rounds of negative selection using 0.1% 5-fluoroorotic acid to eliminate constitutive mutants from the libraries; ~0.001% of cells showed ligand-independent growth after negative selections.

### High-throughput sensor screens

Sensor screens were conducted in duplicate (DSM, shuffle) or quadruplicate (TSM) using 96-well microtiter plates with wells containing 200 µL of solid SD-Trp, -Leu, -Ura, 1.5% agar media (Sigma-Aldrich, USA), and 100 µM test compound. Each well was seeded with approximately 150,000 yeast cells harboring PYR1 variants in *S. cerevisiae* MaV99 (pACT-HAB1), incubated at 30 ºC for three days. Colonies were collected and used in retests with and without the test chemical. Up to 8 validated colonies from each positive well were propagated, and receptor amplicons sequenced. In the DSM and shuffle screens, individual colonies were selected from wells, retested, and Sanger sequenced. To expedite TSM screening, hit validation, and eliminate wells dominated by constitutive mutants, wells with colonies were washed to collect all cells, then retested on control wells lacking test compound; wells that exhibited robust growth without ligand (as evident by a lawn of growth) were eliminated from further consideration. Individual colonies from the remaining wells were subsequently isolated and retested in both the presence and absence of the ligand to confirm ligand-dependent responsiveness. Amplicons for 12 validated clones were sequenced using a modified version of evSeq (*41*) with barcoded primers designed for pBD-PYR1 receptor amplicons. Individual yeast colonies were amplified using GoTaq DNA polymerase (Promega Corporation, USA) and pooled PCR products purified using a Monarch PCR & DNA Cleanup Kit (NEB, USA), and sequenced by Novogene (USA) using paired-end 250 bp reads. Receptor sequences were derived from fastq files after filtering low-quality reads using PYR1_evSEQ (https://github.com/laoda2023/Mutant_Library-Analysis) (*42*). Validated, non-redundant (i.e., unique binding pocket) mutant sensor strains were subsequently tested in dose-response experiments using newly stocked chemicals (DSM and shuffle screens) or compounds from source wells (TSM screen). Screens of targeted receptor variants (TNT, PFAS, coumarins) were conducted similarly, but used ~10^6^ cells plated onto media in 25 cm or 6-well plates. Although this work used MaV99, we have determined that the commercially available strain MaV203 also works. The sequences of the previously described (*43*) pACT-HAB1 and pBD-PYR1 are provided here for reference (https://bit.ly/3EVWOD8; https://bit.ly/4jIg325).

### Receptor library assembly from oligonucleotide pools

Libraries were constructed using the Golden Gate Assembly (GGA) to assemble the PYR1 coding sequence from three ~200-nucleotide blocks PCR amplified from oligonucleotide pools and cloned into pBD-PYR1-156, a domesticated pBD-PYR1 vector with BsaI sites inserted adjacent to codon 50 and with (pBD-PYR1-156-F1; https://bit.ly/453M7J8) or without (pBD-PYR1-156; https://bit.ly/4lUbTFQ) an in-frame stop codon adjacent to the 3’ BsaI site. Oligonucleotides encoding each of the desired mutations were generated using sequence profiles designed in the Targeted-PYR1-Library-Design ShinyApp, which designs the encoded mutation according to user-set parameters, adds flanking *indexset* barcode primers (*44*) to each of the oligonucleotides, and outputs them to a fasta file (https://github.com/weslgeorge/Targeted-PYR1-Library-Design-App). Most assemblies were conducted using oligonucleotides barcoded with block- and library-specific 18-mer primers, allowing PCR amplification of the target blocks from pools designed to harbor multiple libraries. The TSM library did not use barcoded oligonucleotides. It was constructed from three separate Twist oligonucleotide pools, each harboring all designs of single, double, or triple mutations at the target sites for each block. The TSM library was assembled from ten GGA reactions to generate all desired triple-mutant combinations systematically. The final library contained 4.3 × 10^6^ clones; deep sequencing determined that 95.3% of the 324,352 designed mutations were present (see details in **Table S9**). Targeted libraries were assembled combinatorially from oligonucleotide blocks that harbored all desired mutations per block (i.e., all single, double, triple, etc). These were assembled into final screening libraries by running seven GGA reactions that combinatorially swapped the three blocks, each carrying either no mutations (wild type, WT), or all possible single, double, or triple mutations. The seven assemblies were transformed into *E. coli*, and their plasmid was pooled before transformation into MaV99 harboring pACT-HAB1 at ratios relative to the number of potential unique PYR1s per assembly. For TSM library quality control and coverage analysis, library plasmid DNA was diluted to 1 ng/μL, and PYR1-amplicons were amplified using Q5 High-Fidelity 2X master mix (NEB, USA) using TH18-lib primers (**Table S10)**. 1.5 μg of column-purified PCR product was sequenced by Novogene (USA) using 2×250 bp paired-end reads on an Illumina NovaSeq. FastQ files were processed to remove reads with a Q-score lower than 20, and coverage was determined using previously described code (*14*). The theoretical library sizes for the targeted libraries range from 77,327 to 516,647 mutants and were constructed with >10x coverage in both *E. coli* and subsequent *S. cerevisiae* strains for libraries with <100,000 members and >2.5x coverage for libraries with >100,000 members.

To assemble the libraries, we PCR-amplified six oligonucleotide blocks—the three oligonucleotide pool blocks that harbor mutants, and the other three wild-type blocks from a plasmid template. One ng of the oligonucleotide pool templates or WT plasmid DNA was amplified using Q5 High-Fidelity 2X master mix and the primers in **Table S10**. Each reaction included 0.5 µM of each primer and was run for 22 cycles (98 °C 10s, 60 °C 30 s., 72 °C 30 s.; initial denaturation 98 ºC, 30 s.). Amplicons were purified using a Monarch PCR & DNA Cleanup Kit (NEB, USA). ~16 ng of each purified block was then combined with either ~300 ng of the pBD-PYR1-156 or pBD-PYR1-156 bp-F1 (2:1 molar insert-to-vector ratio) to assemble the seven block permutations. BsaI-HFv2 and T4 DNA ligase (NEB, USA) were then added using the manufacturer’s recommendations and cycled (1 minute each 37°C/16°C; 30 cycles). The reaction was transformed into chemically competent Top10; plasmid DNA was used to transform *S. cerevisiae* MaV99 (pACT-HAB1) as described above.

### Design and construction of an improved double-site mutant receptor library (DSM-Hao)

Our initial screens used the previously designed PYR1 DSM library (*14*). This library was built using nicking site saturation mutagenesis(*45*) and harbors many wild-type sequences. We therefore created a new DSM library using GGA from oligonucleotide pools, based on the previous DSM sequence profile with slight modifications; in designing this new library, we sought to reduce the number of constitutive mutants. We, therefore, excluded two strong constitutive single mutants (V83F and F159V) (*46*) from all designs. In addition, evSeq sequencing of a set of constitutive receptors and a separate deep sequencing experiment, conducted to identify mutants depleted from the DSM library after negative selections, together identified 1,188 constitutive double mutants co-located in the same assembly blocks. These double mutants were removed from the designed oligonucleotides, which were synthesized by Twist Bioscience (USA) and used to assemble the library as described above; see **Table S3** for the sequence profile, **Table S4** for the designed oligonucleotides, and **Table S11** for the mutants excluded from the library. The library, created at 99.95% of coverage, harbors 34,657 of the 34,675 designed mutations, as determined by deep sequencing **(Table S9)**.

### Targeted library design using sequence profiles

To create targeted PYR1 libraries with focused small-molecule biosensor targets, sequence profiles were generated using binding pocket mutations from target sensors. Similar molecules in the ligand-receptor dataset were identified using similarity searches and clustering based on atom-pair fingerprints in ChemineR (*21*). The Targeted-PYR1-Library-Design-App (https://github.com/weslgeorge/Targeted-PYR1-Library-Design-App) (*47*) was used to generate a sequence profile for the selected sensors, which were manually curated to eliminate rare mutations and limit the final library sizes to less than 600,000. The final sequence profiles were used to create oligonucleotide pools for assembly using the methods described above. The sequence profiles used for targeted libraries are provided in **Table S3**. The coumarin library sequence profile was constructed from osthole, imperatorin, and isopsoralen sensors. The TNTv1 sequence profile was built from sensors for 2,4-dinitrophenol, chloroxylenol, 2,4,6-trihydroxybenzoic acid, 2,6-dihydroxybenzoic acid, which we selected based on their similarity to TNT; the TNTv2 sequence profile was constructed from the TNT/ADNT/DNT screening hits from TNTv1, to which we added amiprofos-methyl sensor data to increase sequence diversity and the mutations V163W and T162D, which arose fortuitously in some of the TNTv1 sensors. The PFAS library’s sequence profile was built from PFAS sensors identified from screening DSM-Hao (**Table S3**).

### Yeast display titrations

Designs were ordered as eBlocks (IDT, USA) encoding the H60P and N90S mutations for compatibility with yeast display(*48*). Plasmids were assembled using Golden Gate cloning with the yeast display destination vector pND003 (*49*). Designs were confirmed by whole plasmid sequencing (Plasmidsaurus, USA). Plasmids were transformed into *S. cerevisiae* EBY100, and strains were stored at −80 °C using standard methods (*50*). Yeast display was performed essentially as described (*48*) using ligand stocks of 10 mM in DMSO. The final concentration of DMSO for all samples tested, including negative controls, was 2 (v/v)%. Plasmid pJS624 (*14*) encoding a yeast displayed PYR1^mandi/N90S^ was included as a positive control.

### β-galactosidase assays

Sensor strain responses to ligands were investigated using X-gal overlay assays of colonies cultured on test ligand or mock for 3 days. For the quantitative measurements of beta-galactosidase expression in the biosensors, 50 mL cultures of yeast sensor strains were grown overnight then diluted to OD_600_ = 0.02 in SD, -Leu, -Trp liquid media in 1 mL aliquots and varying concentrations of test ligand from 1000X-stocks in DMSO, incubated for approximately 7 - 16 hours then diluted OD_600_ 0.1 in SD-Leu-Trp media and 50 µL of the suspension transferred to opaque flat-bottom Optifine 96-well plates along with 50 µL of Beta-Glo assay mixture (Promega, USA). Luminescence was measured using a Tecan Infinite F200 pro plate reader with shaking (450 rpm for 30 seconds) over 2 hours, using a 200 ms integration time. Relative luminescence (RLU) measurements are plotted using a time point at which the RLU has stabilized (~90 minutes).

## Supplementary Figures

**Figure S1.**
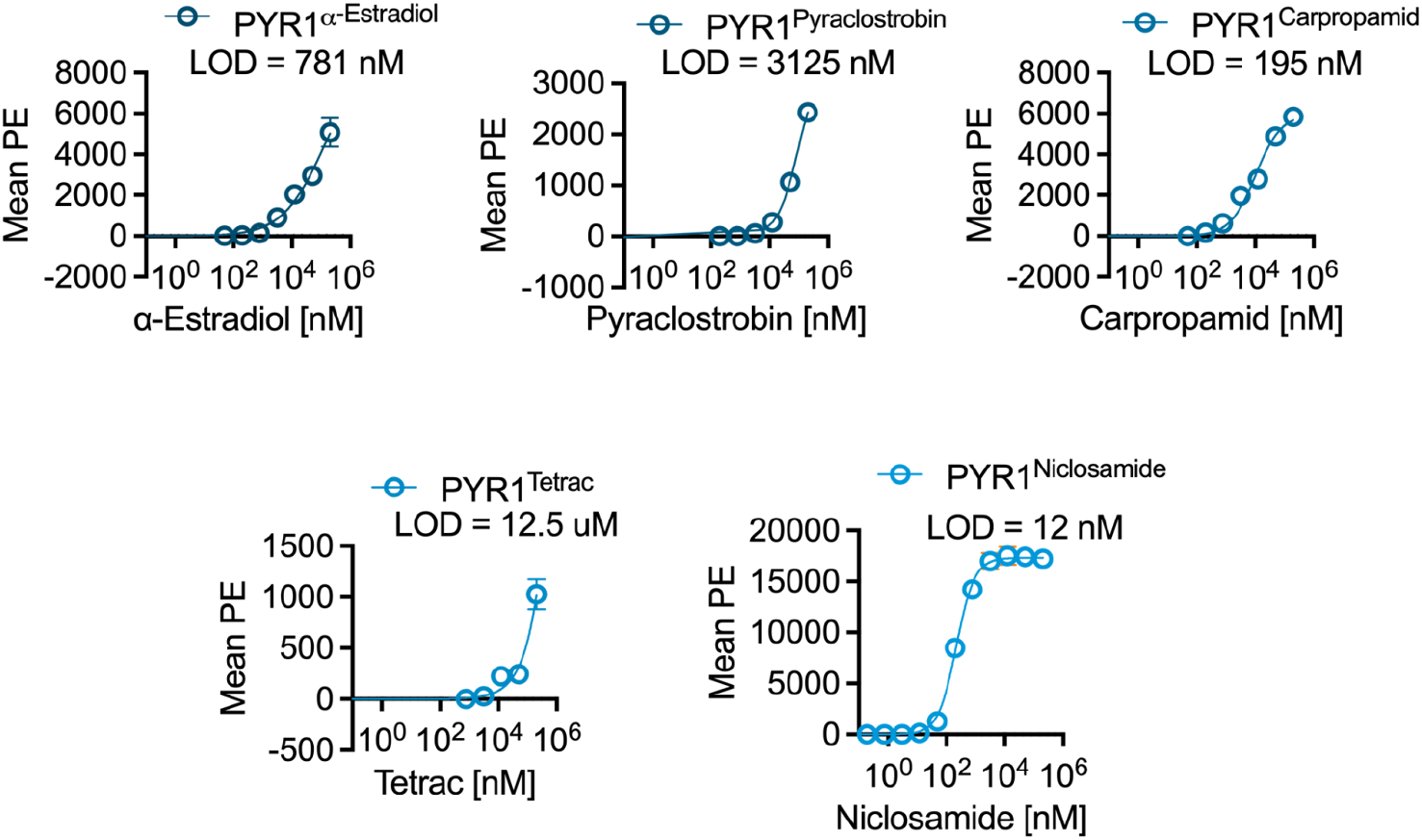
Direct and sensitive binding of PYR1-derived sensors to their target ligands. Dose-response characteristics were quantified for five PYR1 mutants isolated from the primary screens. 17 PYR1 mutants were initially selected; 11 showed detectable binding in the yeast surface display assay, requiring a minimum apparent dissociation constant (K_d_) of 1 µM. The five shown were selected for dose-response validation. Yeast surface displays (YSD) were conducted as previously described (*48*), with the exception that they employed HAB1^T+^, a stabilized variant of HAB1 (*14*). Sensors were prepared for YSD by incorporating H70P and N90S mutations, which are required for proper display of PYR1. Assays were conducted using 200 nM biotinylated HAB1^T+^ and varying target ligand concentrations (380 pM - 200 µM). Error bars are shown as two s.e.m. (n=4; two bio reps, two tech reps). Complex formation is indicated by fluorescence derived from a streptavidin-phycoerythrin conjugate recruited HAB1^T+^ upon ligand binding. The limits of detection (LOD) were calculated using the 3σ method, which is equivalent to the drug concentration that yields a signal equal to 3 times the standard deviation of the blank after subtraction.

**Figure S2.**
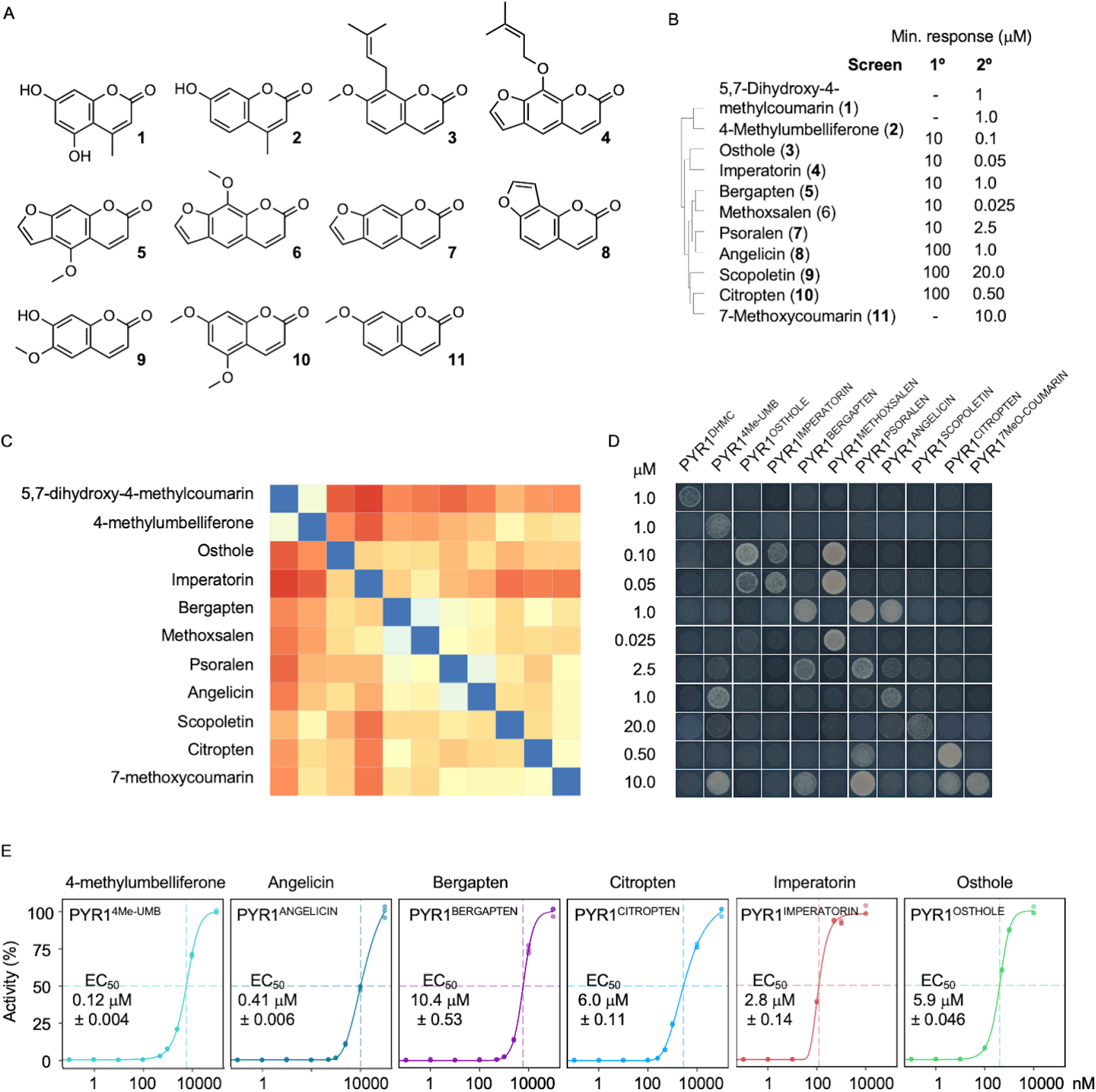
Coumarin-related receptor screening and analysis. (**A**) Structures of the 11 coumarin-related compounds used for PYR1 receptor screening. (**B**) Comparison of sensor sensitivity across screens; all libraries and PYR1 hits are shown in Table S2. Dendogram (left) shows the coumarin chemical similarity based on data in (C). Lack of response in the primary screen (‘-’) indicates that the compounds failed to activate any PYR1 mutant at a concentration of 100 μM. (**C**) Heat map showing pairwise structural similarity between the 11 compounds, based on a distance matrix of pairwise Tanimoto similarity scores, calculated in ChemmineR. (**D**) The cross-reactivity matrix for the most sensitive receptors identified for each ligand, as well as receptor names, is shown above each column. Sensor strains were tested at the on-target minimum activation concentration observed in yeast hybrid growth assays (see panel B). (**E**) Dose-response curves for selected receptors in yeast, quantified using the Beta-Glo assay system. Data points represent individual values (n=3); EC_50_ values with associated standard deviation are shown.

**Figure S3.**
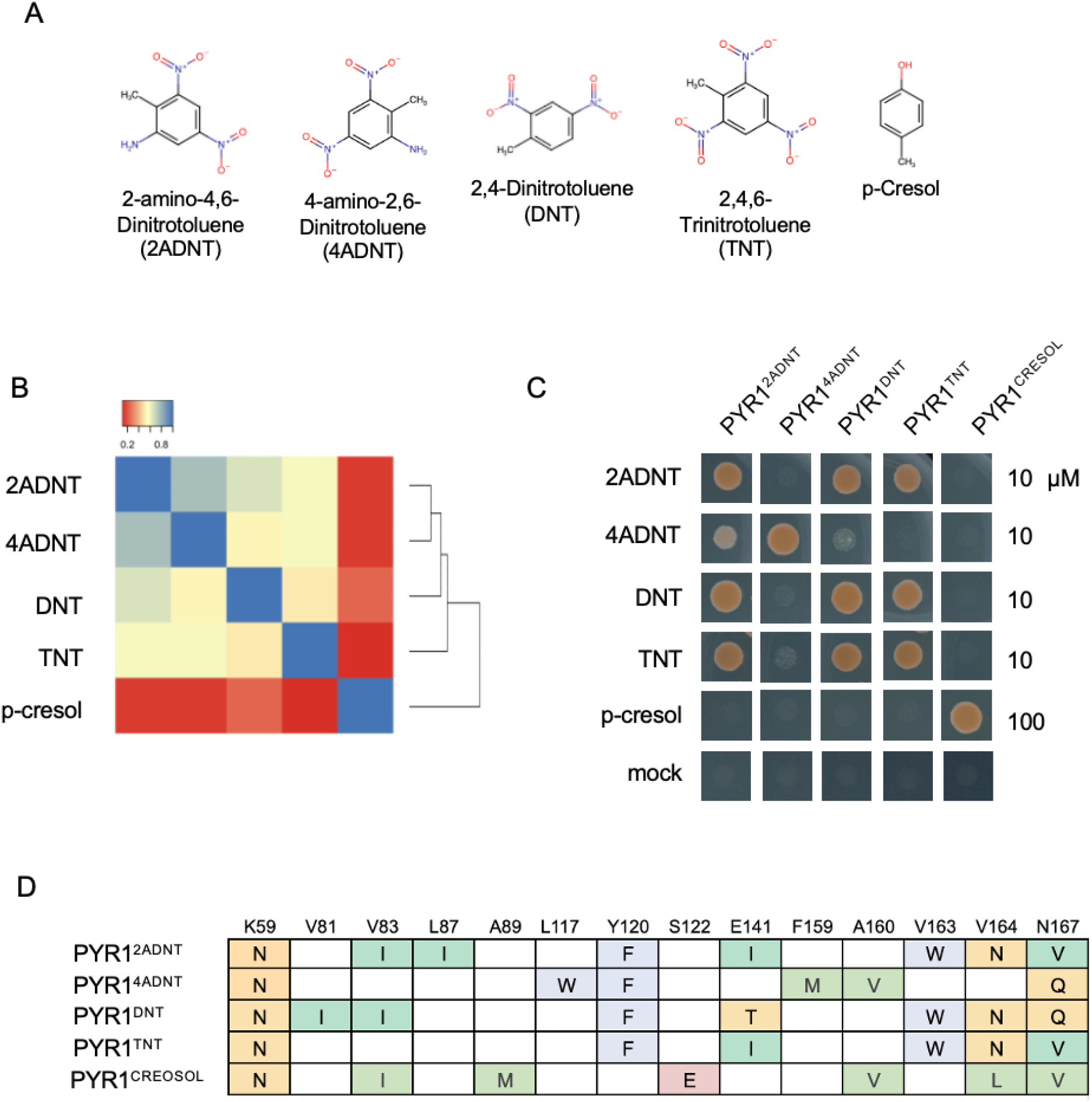
TNT and degradation product sensors. (A) Chemical structures of TNT and related compounds targeted for sensor development. p-Cresol was one of the phenyl aromatics used to generate the input sequence profile and was screened as a positive control. (B) Comparison of the chemical similarity of target molecules by Tanimoto score and (C) The most sensitive sensors isolated from each screen (PYR1^2ADNT^, PYR1^4ADNT^, PYR1^DNT^, PYR1^TNT^, PYR1^CRESOL^) were tested for on- and off-target responses in yeast hybrid growth assays at 10x the minimum concentration required for a growth response. (D) Binding pocket mutations in the sensors.

**Figure S4.**
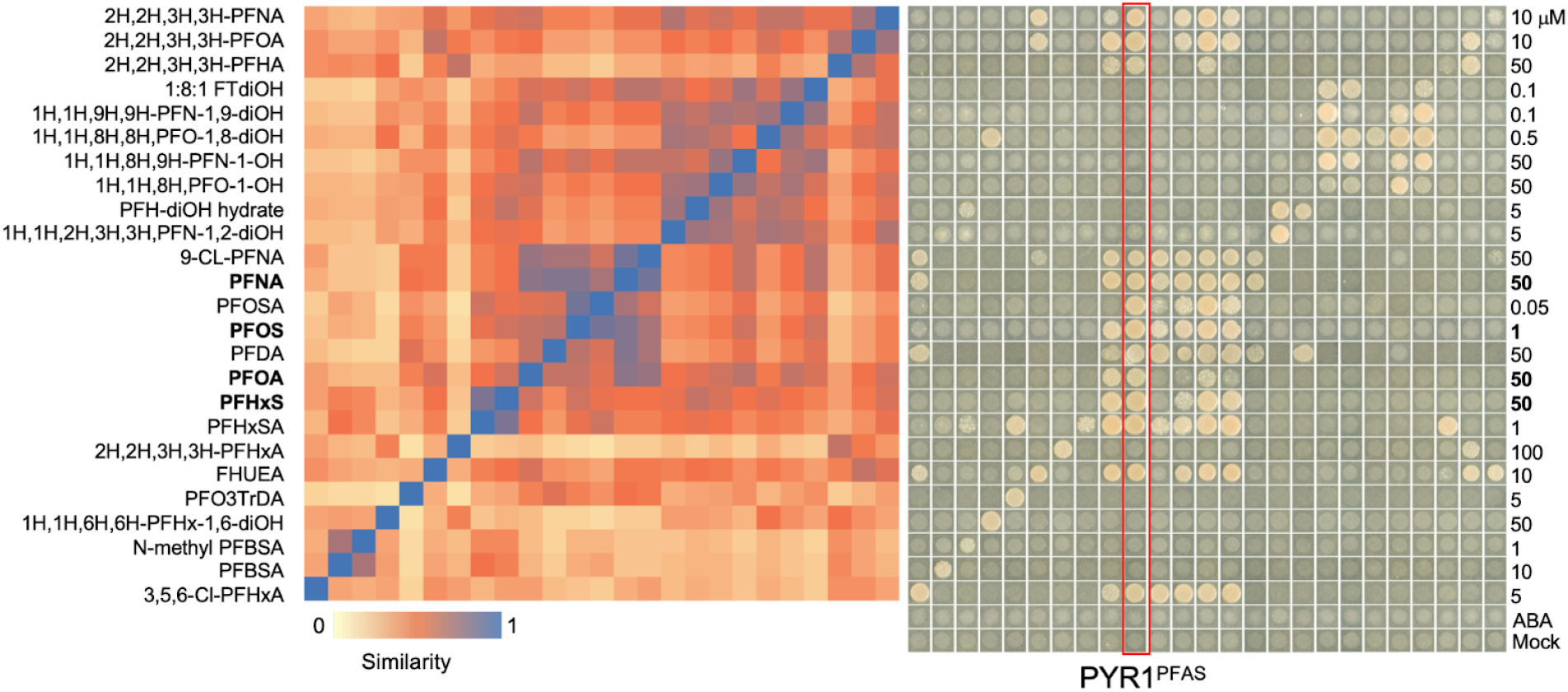
Comparison of the chemical similarity of hit PFAS molecules by Tanimoto score (left) and their on- and off-target responses of their cognate receptor in yeast hybrid growth assays (right). PYR1^PFAS^ was isolated in a screen against PFOA. The strong on- and off-target effects of PFNA, PFOS, PFOA, and PFHxS show that it can be used as a sensor for any of these molecules. Sensor strains were tested at the on-target minimum activation concentration observed in yeast hybrid growth; sensor strains were tested by spotting 2 µL of a single colony suspended in 100 µL TE buffer onto selective media.

